# Chronic FMNL2 overexpression is associated with changes in brain histology and exploratory behavior in adult zebrafish

**DOI:** 10.64898/2026.11.09.765478

**Authors:** Chu Lyu, Huseyin Tayran, Prabesh Bhattarai, Elanur Yilmaz, Caghan Kizil

**Affiliations:** Department of Neurology, Columbia University Irving Medical Center, Columbia University New York, NY, USA; Taub Institute for Research on Alzheimer’s Disease and the Aging Brain, Columbia University Irving Medical Center, Columbia University, New York, NY, USA; New York Genome Center, New York, NY 10013, USA

**Keywords:** FMNL2, Alzheimer’s disease, zebrafish, gliovascular interactions, amyloid-β42, microglia, novel tank diving

## Abstract

Formin-like protein 2 (FMNL2) has been associated with Alzheimer’s disease and cerebrovascular pathology and regulates gliovascular remodeling following amyloid exposure. However, the consequences of sustained FMNL2 elevation remain unclear. We generated a transgenic zebrafish line expressing human FMNL2 under the her4.1 promoter and examined adult brain histology and behavior under baseline conditions and following acute amyloid-β42 (Aβ42) exposure. Histological analyses assessed FMNL2 immunoreactivity, overlap between the glial marker BLBP and endothelial tight-junction marker ZO1, synaptic vesicle protein SV2 labeling, and microglial counts. Exploratory behavior was evaluated using the novel tank test and free-movement pattern Y-maze. Compared with uninjected wild-type controls, uninjected FMNL2-overexpressing fish showed higher FMNL2 immunoreactivity, lower BLBP–ZO1 overlap, fewer normalized SV2 puncta, and higher counts of total and morphologically activated microglia. Acute Aβ42 exposure in wild-type animals produced changes in the same direction across these histological endpoints. No significant additional histological changes were detected following Aβ42 exposure in the overexpression background. In the novel tank test, uninjected FMNL2-overexpressing fish had a higher composite boldness index than uninjected wild-type controls. These findings indicate that sustained FMNL2 overexpression is associated with histological changes resembling several responses to acute Aβ42 exposure, together with altered exploratory behavior. They support further investigation of persistent FMNL2 elevation as a contributor to altered glial-vascular organization and cellular responses relevant to Alzheimer’s disease.

## Introduction

Alzheimer’s disease (AD) is characterized by amyloid-β accumulation and tau pathology, accompanied by synaptic dysfunction, neuroinflammation, and progressive cognitive impairment ^1–4^. However, these processes occur within a broader tissue environment in which cerebrovascular abnormalities can contribute to brain dysfunction. Neuropathological studies demonstrate the frequent coexistence of AD and vascular lesions ^4–13^, while clinical evidence links blood-brain barrier abnormalities to cognitive dysfunction, including changes that occur independently of amyloid and tau biomarkers ^8,14–19^. Understanding how vascular and neurodegenerative processes intersect is therefore important for identifying mechanisms that influence the development and progression of disease.

Interactions between astroglial cells and the cerebral vasculature contribute to the organization and maintenance of the brain’s vascular interface ^20,21^. Changes in these interactions may influence tissue homeostasis, inflammatory responses, and the handling of extracellular material. Remodeling of the glial-vascular interface may form part of a response to pathological stimuli, but the consequences of maintaining an altered organization over time remain less clear ^14–16,22,23^. Identifying the molecular regulators of these interactions may help explain how an initially responsive cellular process becomes persistently altered during disease.

Our previous work identified Formin-like protein 2 (FMNL2) through a gene-by-cardiovascular-risk-factor analysis of AD and linked elevated FMNL2 expression to cerebrovascular and AD-associated pathology ^24^. Higher brain FMNL2 expression was associated with infarcts and with amyloid and phosphorylated tau deposition, with prominent expression in astroglial cells ^24,25^. FMNL2 belongs to the formin family of actin regulators and promotes actin filament elongation and cellular protrusion ^26–28^. These functions provide a basis for investigating its role in the structural organization and remodeling of glial-vascular interactions.

Experimental findings further support a role for FMNL2 in the response to amyloid. In adult zebrafish, acute Aβ42 exposure increased astroglial *fmnl2a* expression and was accompanied by reduced association between astroglial endfeet and blood vessels ^24^. Knockdown of *fmnl2a* prevented this remodeling, reduced microglial activation, and increased amyloid deposition. These observations implicated FMNL2-dependent remodeling in the tissue response to amyloid and suggested that elevated FMNL2 expression may participate in an adaptive response rather than simply mark pathological damage. However, they did not establish the consequences of sustained FMNL2 elevation in the absence of an acute amyloid stimulus.

This distinction is important because persistent expression of a remodeling regulator may have different consequences from its transient recruitment during an acute response. Sustained FMNL2 elevation could maintain changes in glial-vascular organization or influence associated synaptic and immune responses even without an additional pathological challenge. Alternatively, it could modify the response to subsequent amyloid exposure. Whether elevated FMNL2 alone is sufficient to produce such changes, and whether it alters the effects of acute Aβ42 exposure, remain unresolved.

To address these questions, we generated a transgenic zebrafish line expressing human FMNL2 ^29^ under the her4.1 promoter ^30,31^. We examined adult animals under untreated baseline conditions and following cerebroventricular injection of PBS or Aβ42. Histological analyses assessed FMNL2 immunoreactivity, overlap between the glial marker BLBP and tight-junction marker ZO1, SV2 labeling, and total and morphologically activated microglial counts. We also examined novel-tank exploration and spontaneous turning strategies in the free-movement pattern Y-maze. This approach allowed us to investigate the consequences of sustained FMNL2 overexpression and determine whether the measured histological and behavioral outcomes were further altered by acute amyloid exposure.

## Materials and Methods

### Animal maintenance and ethics declaration

Zebrafish (*Danio rerio*) of AB wild-type strain and FMNL2 Overexpression line of both male and female were bred in-house and raised in the Columbia University Animal Care Facility. Fish were housed in groups of 20-25 per 2.8 L tank on a re-circulating system (Aquaneering, CA, USA). The system uses reverse osmosis water adjusted with conductivity (range: 500–1,000 µS/cm) and pH (∼7.4). Room and water temperatures were kept at around 27°C on a consistent 14h-light and10h-dark cycle. Fish were fed fry food from 5 days post-fertilization until adulthood when the diet changed into live three times/day (once/day on weekends).

6- to 12- month-old transgenic fish lines and their negative siblings of both sexes were used. Animals from the same clutch were randomly selected for each experimental setup. For zebrafish, all animal experiments were performed in accordance with the applicable regulations and approved by the Institutional Animal Care and Use Committee (IACUC) at Columbia University (protocol number AC-AABN3554). Animals from the same fish clutch were randomly distributed for each experimental group. Animals were handled with caution to reduce suffering and overall animal numbers.

### Generation of Human FMNL2 overexpression line

#### Vector Construction for Tol2 Transgenesis

Human FMNL2 cDNA (kindly provided by Dr. John Copeland ^29^) was amplified via PCR using forward and reverse primers containing FseI and AgeI restriction sites at their 5 ‘ ends, respectively. The PCR reaction was performed using Q5 High-Fidelity DNA Polymerase (NEB, Cat. No. M0491S) under the following conditions: an initial denaturation step at 98 °C for 5 min, followed by 40 cycles of denaturation at 98 °C for 20 s, annealing at 63 °C for 15 s, and extension at 72 °C for 2 min and a final extension at 72 C for 10 min. The resulting PCR product was purified using the GeneJET PCR Purification Kit (Thermo Scientific, Cat. No. K0702). For cloning, 1 ug of the pTol(her4.1:T2A_GFP_PolyA) plasmid backbone and 500 ng of the purified FMNL2 PCR fragment were digested with FseI and AgeI-HF restriction enzymes (NEB, Cat. No. R0588S and R3552S, respectively) at 37 °C for 1 h. Digested products were then run on 1% agarose gel at 100 V for 50 min, excised, and subsequently purified using the GeneJET Gel Extraction Kit (Thermo Scientific, Cat. No. K0692). 200 ng of the digested backbone and 125 ng of the insert (Human FMNL2 gene) were ligated overnight at 4 °C using T4 DNA Ligase (NEB, Cat. No. M0202S) to yield the final pTol(her4.1:Human FMNL2 _T2A_GFP_PolyA) construct. The ligation mixture was transformed into competent E. coli DH5α cells. Following a 1 h recovery at 37 °C, 100 μl of the bacterial culture was plated onto LB agar containing ampicillin (100 μg/ml) and incubated overnight at 37 °C. Individual colonies were selected and inoculated into liquid LB medium supplemented with 100 μg/m ampicillin and were grown overnight at 37 °C with shaking at 250 rpm. Plasmid DNA was isolated from overnight cultures using the GeneJET Plasmid Miniprep Kit (Thermo Scientific, Cat. No. K0503). The final plasmid was then confirmed by Sanger sequencing.

#### Microinjection into Single-Cells Embryos

The FMNL2 overexpression line was established by microinjecting a 2 nl mixture containing pTol(her4.1:Human FMNL2 _T2A_GFP-PolyA) (7.5 ng/µl) and Tol2 transposase mRNA (12.5 ng/µl) into one-cell stage WT-AB zebrafish embryos. Injected F0 mosaic embryos were delivered to fish room and raised to adulthood.

#### Screening F0 Founders and Establishing F2 Stable Lines

Potential F0 adult fish were outcrossed to WT-AB fish to identify germline-transmitting founders.30 embryos of 2-3 days post-fertilization from each cross were pooled into separate PCR tubes and their genomic DNAs were extracted by alkaline lysis method ^32^. 300 ul from 50 mM NaOH was put into each tube and tubes were heated at 95 C for 20 min to release the genomic DNAs. Following, all tubes were kept on ice to cool to 4 C and then 30 ul from 1 M Tris-HCI (pH:8.0) was added into tubes to neutralize the basic solution. All tubes were then centrifuged to pellet the debris for 5 min at 10.000 rpm and supernatants were taken into new tubes. 3 ul from these supernatants was used in the following 25-ul PCR reactions with Dream Taq DNA polymerase (Thermo Scientific, EP0705) with the conditions : an initial denaturation at 95 C for 5 min, followed by 40 cycles of denaturation at 95 C for 30 sec, annealing at 63 C for 20 sec, extension at 72 for 1 min 30 sec and a final extension at 72 C for 10 min. 15 microliters from each PCR reaction were then loaded into 1% agarose gel and run for 40 min at 100V. T2A-GFP part of the construct was screened with the following primers: forward primer : 5’-CAATGACCGGTAGCGGAGAGGGCAGAGGAAGTCTTCTA - 3’ and reverse primer : 5’-AGAGGCTCGAGTTTACTTGTACAGCTCGTCCATGCCG - 3’ The progenies from 5 confirmed F0 founders were grown to adulthood to establish the stable F1 adults. This outcrossing and screening process was repeated and tracked by GFP expression and confirmed by PCR to obtain the subsequent F2 generation.

### Tissue processing and analyses

6-month-old transgenic animals and their negative siblings of both sexes with no treatment and 8-month-old groups were injected with PBS and Amyloid-βeta42 (Aβ42) in the cerebral ventricle by cerebroventricular microinjection (CVMI). Animals were euthanized and processed for histological experiments at 3 days post-injection. The zebrafish heads were first dissected and fixed in 2% paraformaldehyde at 4°C overnight, then decalcified in 20% Sucrose with 20% ethylenediaminetetraacetic acid (EDTA) solution for 2 days. On day 3, fish heads were embedded in O.C.T., cryosectioned, and stored as 12-µm-thick slices on glass slides at −20°C. Sections were warmed to room temperature, washed three times with 0.3% PBSTx (PBS with 0.03% Triton X-100), and incubated with primary antibodies overnight at 4°C. For antigen retrieval, slides were heated to 85 °C in 10mM sodium citrate (pH 6.0) for 15 minutes prior to applying primary antibodies. The next day, sections were washed three times with 0.3% PBSTx, incubated with corresponding secondary antibodies and DAPI for 2 hours at room temperature, washed with 0.3% PBSTx, and mounted with 70% glycerol. The following primary antibodies were used: mouse anti-FMNL2 (Proteintech, Cat. No. 68551-1-Ig), mouse anti-ZO-1 (Thermo Fisher Cat. No. 33-9100), mouse anti-SV2A (DSHB Cat. No. SV2), rabbit anti-l-plastin (gift from Belgium), rabbit anti-BLBP (Abcam, Cat. No. ab279649).

### Imaging and quantification

Images were acquired using a Zeiss confocal LSM800 microscope under 20X objective. 6 to 12 histological sections within the telencephalon were used per animal for the quantitative analysis (n = 4 animals in every experimental group, 2 for each sex). All Images were quantified using Fiji software (version 2.0.2) with automated macro code for batch processing for each set of staining combinations. Output files from ImageJ were organized and further analyzed using Python (v3.10.7; Supplementary File 1). FMNL2 and BLBP channels were first colocalized to identify FMNL2-expressing glial cells. FMNL2 expression within these cells was quantified as the total FMNL2 fluorescence intensity divided by the total FMNL2-positive area. For analysis of BLBP and ZO1 overlap, a region of interest (ROI) along the ventrolateral region where BLBP and ZO1 meet was selected and standardized to the same area across all samples. Within this region, BLBP and ZO1 were colocalized to quantify their overlapping area, which was presented as a percentage of the total ROI area. The same anatomical region and standardized area were used for SV2 analysis. A Difference of Gaussians filter was applied to the SV2 channel to reduce background noise and facilitate puncta detection using the Analyze Particles function. DAPI-positive nuclei within the same ROI were quantified using the same function, and SV2-positive puncta counts were normalized to DAPI counts. Microglia were classified into activated, intermediate, and resting morphological categories based on their morphology and branching complexity. L-plastin-stained microglia were segmented using local enhancement, thresholding, and particle analysis, with individual cells extracted as regions of interest (ROIs). For each cell, morphological features, including area, perimeter, circularity, solidity, and feret diameter, as well as branching features obtained through skeletonization (number of branches, junctions, endpoints, average branch length, and total branch length), were quantified. All data were combined together for subsequent analysis. Six main morphological features were used for clustering: circularity, solidity, branches, endpoints, junctions, total branch length. Features were standardized using z-score normalization and subjected to principal component analysis (PCA) for dimensionality reduction. The first two principal components were used for K-means clustering (k=3). Based on cluster-level morphological characteristics, microglia were assigned to three categories: activated (less ramified, more rounded morphology), intermediate, and resting (highly ramified morphology). Cluster separation was evaluated using the silhouette score, and the proportion of each morphological category was calculated for individual images and fish (Tables S1 and S2).

### Behavior experiment

Two different batches of wildtype were used in the experiment to ensure reproducibility across independent cohorts. One batch of n = 16 per group was used in wildtype comparison while another batch wildtype of n = 16 was used in comparison to overexpression group. However, two fish in this wildtype group froze for the entirety or majority of experiment and were thus removed. n = 12 was used per overexpression group. All groups had roughly 50:50 male/female ratio. Both Novel Tank Diving Test (NTT) and Y-maze experiments were completed between 1 pm to 5pm and controlled by a laptop using the Zantiks AD automated tracking system (Zantiks Ltd., Cambridge, UK), following established protocols. Fish first went through NTT and then Y-maze test. Water was changed in between each test for each fish. Files containing the behavioral data and video recording were exported for subsequent analysis, which was done using custom Python scripts (v3.10.7; Supplementary File 2). For novel tank diving test, individual fish were placed in a transparent tank (140 mm W, 200 mm L and 150 mm H) with 1.5L of aquarium water for 300s (Fig. S2). The x and y coordinates of fish were recorded per 20 milliseconds and were used for subsequential analysis. A composite boldness index was calculated using min-max-normalized distance from the bottom and total distance traveled, weighted at 2/3 and 1/3, respectively: Boldness index = 2/3 × normalized distance from the bottom + 1/3 × normalized total distance traveled ^133^. Mean distance from the bottom was calculated using the difference between the maximum y coordinate (140 mm) and each recorded Y-coordinate, averaged across all tracking points. Total distance traveled was calculated as the sum of Euclidean distances between consecutive x,y coordinates. Spatial occupancy heatmaps were generated using two-dimensional histograms of the mean of the recorded x,y coordinates. Both groups were plotted using identical spatial ranges, bin numbers (60 × 60), and occupancy scales to allow direct comparison of swimming distribution between groups. Swimming speed was calculated for each 10-s interval by dividing the total distance traveled during that interval by 10s. For Y-maze test, in one system, a set of two fish was placed separately into two identical acrylic Y-arena insert (three identical arms with the dimensions of H140 x L50 x W19 mm) with 3L of aquarium water for one hour (Fig. S3). Fish movements were documented as they entered and exited each arm and turning choices were assessed. Transitions between the three arms of the Y-maze were classified as left (L) or right (R) turns based on the direction of movement between consecutive arms. Sequential turning patterns were analyzed by grouping consecutive L/R turns into four-turn sequences (tetragrams). All 16 possible tetragram patterns were identified, and their frequencies were calculated as percentages of the total number of tetragrams for each fish. Overlapping sequences were included, such that each successive tetragram shared three turns with the preceding sequence.

### Statistical analysis

All statistical analyses were performed in GraphPad Prism (version 11.1.0). The most appropriate statistical test for each data set was determined by its distribution. For comparison of multiple experimental groups, the appropriate post-hoc multiple comparison test was applied depending on the primary test used (e.g., Tukey’s test following an ordinary one-way ANOVA). Error bars shown in graphs are the mean with SEM values. P values are indicated using GP style: 0.1234 (ns), 0.0332 (*), 0.0021 (**), 0.0002 (***), <.0001 (****). Damaged sections accounted for less than 1% of histological samples and were excluded from analysis. Behavioral exclusions are described above.

## Results

### Astrocytic FMNL2 overexpression reduces synaptic puncta, alters astrocyte coverage of blood vessels and increases the number of microglia in the adult zebrafish brain

To investigate the consequences of sustained FMNL2, we generated a zebrafish line expressing the human FMNL2 coding sequence under the her4.1 promoter using Tol2-mediated transgenesis (**Fig. 1A**). The transgenic line followed Mendelian inheritance, remained genetically stable across generations, and produced viable and fertile progeny. Heterozygous transgenic animals were used to examine the effects of elevated human FMNL2 expression in the adult brain. Immunofluorescence staining confirmed higher FMNL2 immunoreactivity in transgenic animals than in wild-type controls (**Fig. 1B**; Nested t-test, t(6) = 6.19, P < 0.001). We next examined whether sustained FMNL2 elevation was accompanied by changes in histological markers associated with gliovascular organization, synapses, and microglial responses. We assessed overlap between the glial marker BLBP and the tight-junction marker ZO1, quantified the synaptic vesicle marker SV2, and evaluated L-plastin-positive microglial cells and their morphology (**Fig. 1C, D**). Compared with wild-type controls, FMNL2-overexpressing animals showed reduced BLBP-ZO1 overlap (**Fig. 1C**; Nested t-test, t(6) = 6.24, P < 0.001) and lower normalized SV2 counts (**Fig. 1D’**; Nested t-test, t(6) = 3.25, P = 0.018). Total microglial counts were higher in the overexpression group (Welch’s t-test, p = 0.0473), and counts of cells classified morphologically as activated microglia were also increased (**Fig. 1D”**; Nested t-test, t(6) = 4.19, P = 0.006; Nested t-test, t(6) = 3.08, P = 0.022). Together, these findings indicate that sustained FMNL2 overexpression is associated with reduced astrocyte and vascular overlap and SV2 labeling, accompanied by increased total and morphologically activated microglial counts in the adult zebrafish brain.

**Figure 1.**
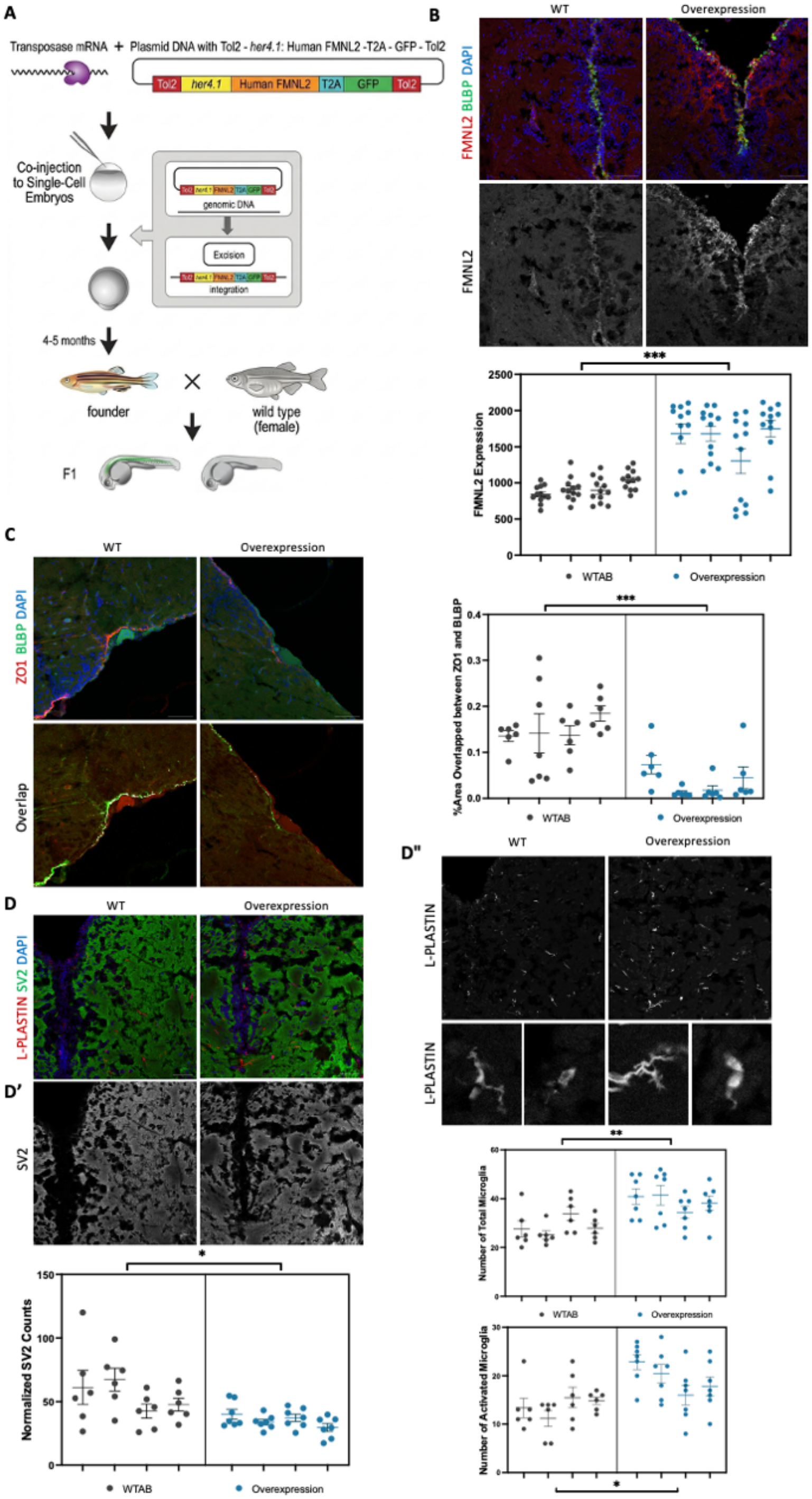
Chronic FMNL2 overexpression is associated with altered BLBP-ZO1 overlap, SV2 labeling, and microglial counts in adult zebrafish. **(A)** Schematic of Tol2-mediated generation of the her4.1-driven human FMNL2-T2A-GFP transgenic line, including injection into one-cell-stage embryos and outcrossing of founder animals. **(B)** Representative brain sections from wild-type (WT) and FMNL2-overexpressing (OE) animals stained for FMNL2 (red), BLBP (green), and DAPI (blue). The FMNL2 channel is shown separately in grayscale below. **(C)** Representative sections stained for ZO1 (red), BLBP (green), and DAPI (blue). Lower images show the ZO1 and BLBP channels with their overlap highlighted in white. **(D)** Representative sections stained for L-plastin (red), SV2 (green), and DAPI (blue), with individual SV2 and L-plastin channels shown in grayscale. Bottom images illustrate cells classified morphologically as resting or activated microglia in each group. **(E)** Quantification of normalized FMNL2 immunoreactivity, BLBP-ZO1 overlap, normalized SV2 counts, total microglial counts, and activated microglial counts. Bars show means with SEM; points represent one measurement per brain section within telencephalon region. Histological sampling comprised four animals per group, two of each sex, with 6-12 sections analyzed per animal. Statistical comparisons used two-tailed nested t-test. *p < 0.05, ***p < 0.001, ****p < 0.0001. Scale bars, 50 μm.

### FMNL2 overexpression produces histological changes resembling those induced by acute Aβ42 exposure

To determine whether chronic FMNL2 overexpression alters the response to acute amyloid exposure, we examined FMNL2 immunoreactivity, BLBP-ZO1 overlap, SV2 labeling, and microglial counts in wild-type and FMNL2-overexpressing animals following cerebroventricular injection of PBS or Aβ42. PBS-injected animals served as vehicle controls.

We first assessed the response to Aβ42 in wild-type animals. Compared with WT PBS controls, WT Aβ42 animals showed increased FMNL2 immunoreactivity, reduced BLBP-ZO1 overlap, lower normalized SV2 counts, and increased total and activated microglial counts (**Fig. 2A-C**). We then examined whether FMNL2-overexpressing animals displayed similar changes in the absence of an acute Aβ42 challenge and whether additional Aβ42 exposure altered these outcomes. FMNL2 immunoreactivity differed among the four groups (**Fig. 2A**; Nested one-way ANOVA, F(3, 12) = 10.56, P = 0.0011). Expression was higher in WT Aβ42 animals than in WT PBS controls (Tukey’s adjusted P = 0.0384). Both OE PBS and OE Aβ42 animals also showed higher FMNL2 immunoreactivity than WT PBS controls (adjusted P = 0.0084 and P = 0.0008, respectively). No significant differences were detected between WT Aβ42 and OE PBS animals (adjusted P = 0.8204), WT Aβ42 and OE Aβ42 animals (adjusted P = 0.1500), or OE PBS and OE Aβ42 animals (adjusted P = 0.4995). BLBP-ZO1 overlap also significantly differed between WT PBS control and three groups (WT Aβ42, OE PBS, and OE Aβ42) (**Fig. 2B**). Three groups (WT Aβ42, OE PBS, and OE Aβ42) all showed significantly lower overlap compared to the WT PBS control. Their adjusted p-values are all statistically significant (0.0043), (0.0112), and (0.0101), respectively. The remaining comparisons between those three altered groups are highly non-significant (p = 0.9535), (p = 0.9617), and (p > 0.9999), which means that BLBP-ZO1 overlap doesn’t differ between these three groups. Normalized SV2 counts differed significantly among groups (**Fig. 2C’**; Nested one-way ANOVA, F(3, 12) = 7.022, p = 0.0056). Counts were lower in WT Aβ42, OE PBS, and OE Aβ42 animals than in WT PBS controls (Tukey’s adjusted p = 0.0176, 0.0275, and 0.0080, respectively). No significant differences were detected between WT Aβ42 and OE PBS animals (adjusted p = 0.9943), WT Aβ42 and OE Aβ42 animals (adjusted p = 0.9485), or OE PBS and OE Aβ42 animals (adjusted p = 0.8628).

**Figure 2.**
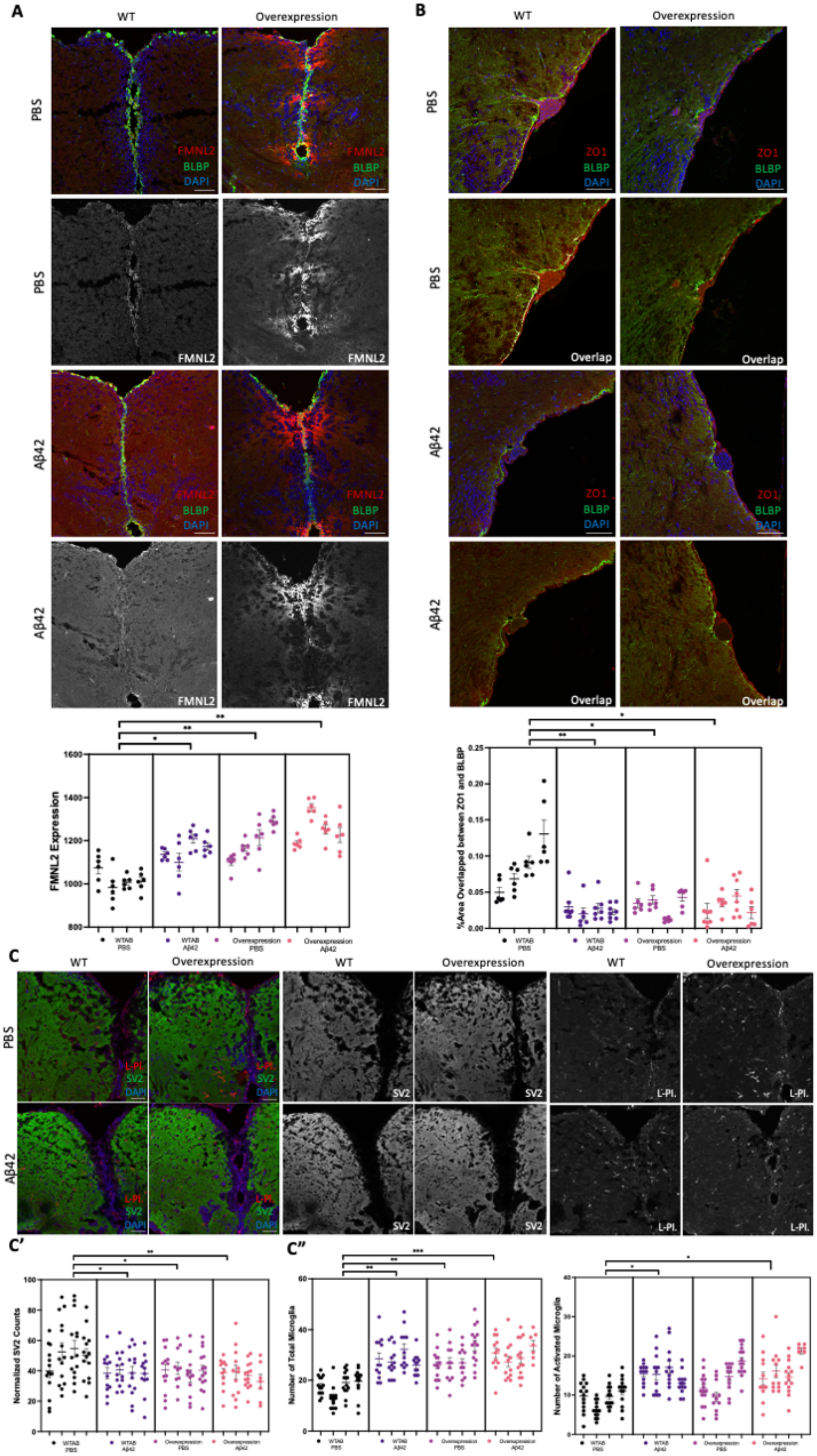
Chronic FMNL2 overexpression and acute Aβ42 exposure are associated with overlapping histological changes. Wild-type (WT) and FMNL2-overexpressing (OE) animals received cerebroventricular injections of PBS or Aβ42, and brains were collected three days after injection. (**A**) Representative sections stained for FMNL2 (red), BLBP (green), and DAPI (blue). Individual FMNL2 channels are shown in grayscale beneath the corresponding merged images. **(B)** Representative sections stained for ZO1 (red), BLBP (green), and DAPI (blue). Corresponding images below show the ZO1 and BLBP channels with their overlap highlighted in white. (**C**) Representative sections stained for L-plastin (red), SV2 (green), and DAPI (blue), with individual SV2 and L-plastin channels shown in grayscale. (**D**) Quantification of normalized FMNL2 immunoreactivity, BLBP-ZO1 overlap, normalized SV2 counts, total microglial counts, and activated microglial counts across WT PBS, WT Aβ42, OE PBS, and OE Aβ42 groups. Bars show means with SEM; points represent one measurement per brain section within telencephalon region. Histological sampling comprised four animals per group, two of each sex, with 6-12 sections analyzed per animal. Statistical comparisons used nested one-way ANOVA followed by Tukey’s multiple-comparisons test. Adjusted P values are reported for pairwise comparisons. *adjusted p < 0.05, **adjusted p < 0.01, ***adjusted p < 0.001, ****adjusted p < 0.0001. Scale bars, 50 μm.

Total microglial counts differed significantly among groups (**Fig. 2C”**; Nested one-way ANOVA, F(3, 12) = 14.60, p = 0.0003). Counts were higher in WT Aβ42, OE PBS, and OE Aβ42 animals than in WT PBS controls (Tukey’s adjusted p = 0.0012, 0.0012, and 0.0005, respectively). No significant differences were detected between WT Aβ42 and OE PBS animals (adjusted p > 0.9999), WT Aβ42 and OE Aβ42 animals (adjusted p = 0.9073), or OE PBS and OE Aβ42 animals (adjusted p = 0.8957). Activated microglial counts also differed significantly among groups (**Fig. 2C”**; Nested one-way ANOVA, F(3, 12) = 4.921, p = 0.0187). Counts were higher in WT Aβ42 and OE Aβ42 animals than in WT PBS controls (Tukey’s adjusted p = 0.0483 and p = 0.0175, respectively), whereas the difference between WT PBS and OE PBS animals was not significant (adjusted p = 0.1640). No significant differences were detected among WT Aβ42, OE PBS, and OE Aβ42 animals (all adjusted p > 0.5).

Thus, chronic FMNL2 overexpression in PBS-injected animals was associated with changes in the same direction as those observed following acute Aβ42 exposure in wild-type animals across the measured histological endpoints. Additional Aβ42 exposure produced no statistically detectable change in these endpoints within the OE background under the conditions examined..

### Chronic FMNL2 overexpression alters behavioral responses in adult zebrafish

The histological analyses indicated that chronic FMNL2 overexpression was associated with several changes also observed following acute Aβ42 exposure in wild-type animals. We therefore asked whether FMNL2 overexpression alone was accompanied by behavioral differences and whether additional Aβ42 exposure altered behavioral outcomes in the OE background. We used the novel tank test to assess vertical exploration and locomotor activity and Y-maze to examine spontaneous turning strategies, over time.

### Chronic overexpression of FMNL2 changes boldness in NTT

The NTT assesses the instinctive behavior of zebrafish in a novel environment and provides multiple readouts^34^. We first compared WT fish with Aβ42-injected WT fish to establish that the acute amyloid-β challenge showed a different response in this assay. We focused on measures that best reflected risk assessment ability because of its relevance in AD patients ^35–37^. This particular trait was assessed using a composite boldness index incorporating distance from the bottom and total distance traveled, with greater distance from the bottom providing the primary measure of upper-tank exploration. We also examined swimming speed and the spatial distribution of fish throughout the tank to further characterize locomotor activity and tank exploration. WT animals exposed to Aβ42 showed a higher boldness index than WT animals (**Fig. 3A**; Welch’s *t*-test, *t*(30) = 2.04, *p* = 0.050). OE animals also had a higher boldness index than WT animals (**Fig. 3D**; Mann-Whitney test, *U* = 45, *p* = 0.045). In contrast, Aβ42 exposure did not further alter the boldness index in FMNL2-overexpressing animals (**Fig. 3G**; Mann-Whitney test, *U* = 69.5, *p* = 0.899). Analysis of swimming speed over the 300-s recording period revealed time-specific differences between WT and WT Aβ42 animals. Repeated-measures two-way ANOVA showed a significant main effect of time (Geisser-Greenhouse-corrected *F*(5.260, 157.8) = 2.539, *p* = 0.0282). Šídák’s multiple-comparisons test identified higher swimming speed in WT Aβ42 animals at 30, 50, 70, 150, 160, 180, and 190 s (**Fig. 3B**; adjusted *p* < 0.05). Comparison of WT and OE revealed that OE animals swam faster at 220, 260, and 280s (**Fig. 3E**; Šídák’s adjusted p = 0.0217, 0.0166, and 0.0435, respectively). No significant differences in swimming speed were detected between OE and OE Aβ42 animals at any time point (**Fig. 3H**). Spatial occupancy heatmaps provided a qualitative visualization of tank use, with WT Aβ42 and OE animals appearing to spend more time in the upper region of the tank (**Fig. 3C,F,I**). Together, these findings suggest that chronic FMNL2 overexpression was associated with an altered boldness-related behavioral profile, while additional Aβ42 exposure produced no detectable further change in the measured boldness or swimming-speed outcomes in the OE background.

**Figure 3.**
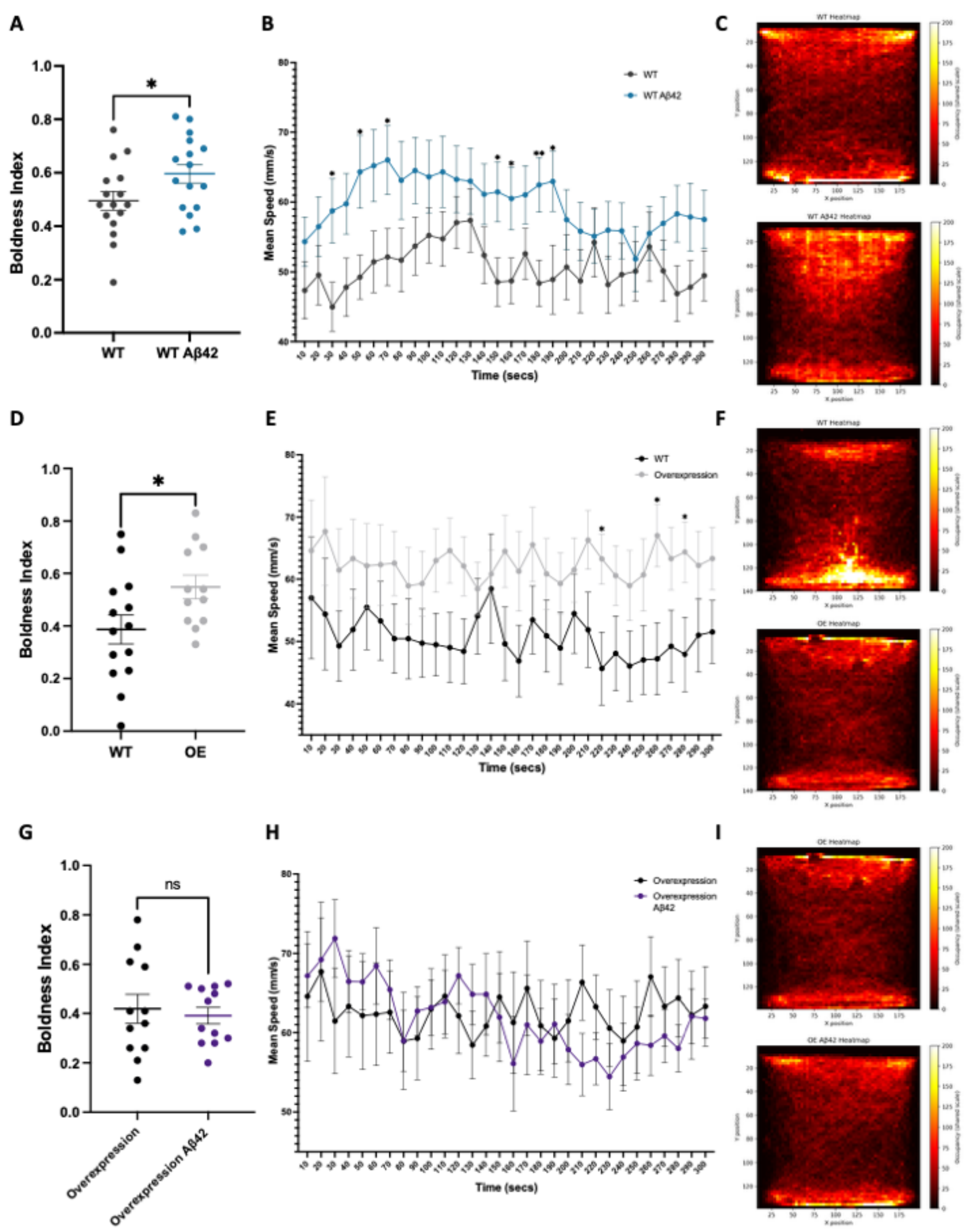
Novel tank exploration and swimming speed in wild-type and FMNL2-overexpressing zebrafish. (**A–C**) Uninjected WT versus Aβ42-injected WT fish. (**D–F**) Uninjected WT versus uninjected OE fish. (**G–I**) Uninjected OE versus Aβ42-injected OE fish. Panels A, D, and G show the composite boldness index, calculated from normalized mean distance from the tank bottom and total distance traveled, weighted by two-thirds and one-third, respectively. Panels B, E, and H show swimming speed in successive 10-s intervals during the 300-s test. Panels C, F, and I show spatial heatmaps on a common color scale within each comparison. Color indicates the mean fraction of recording time fish spent in each spatial bin. Bars and trajectories show means with SEM; points in the boldness plots represent individual fish. Sample sizes were n = 16 for WT vs n = 16 for WT Aβ42, n = 14 for WT vs n = 12 for OE, n = 12 for OE vs n = 12 for OE Aβ42. Boldness indexes were compared using Welch’s t-test for WT baseline versus WT Aβ42 and Mann-Whitney tests for the other two comparisons. Swimming-speed trajectories were analyzed using repeated-measures two-way ANOVA followed by Šídák’s multiple-comparisons tests at individual time points. Asterisks above trajectories indicate between-group differences at the indicated time points. ns, not significant; *p < 0.05, **p < 0.01; time-point comparisons use multiplicity-adjusted p values.

### Repetition and alternation were common turning strategies across groups in the Y-maze

We examined spontaneous turning strategies in the free-movement pattern Y-maze ^38,39^. For each fish, we calculated the frequencies of the 16 possible four-turn sequences (**Fig. 4A, C, E**) and combined complementary left- and right-turn sequences into eight pairs (**Fig. 4B, D, F**). If successive turns are independent and left and right turns are equally likely, the expected frequency is 6.25% for each sequence and 12.5% for each pair. Across all groups, the observed mean frequencies of repetition (LLLL/RRRR) and alternation (LRLR/RLRL) both exceeded the 12.5% reference value. Although repetition and alternation remained the predominant strategies across all groups, the degree of separation among the turning strategies differed across experimental conditions. In WT baseline animals, relatively few differences were observed among the paired strategies, with the predominant repetition and alternation strategies showing limited separation from the remaining patterns. Following Aβ42 exposure, WT animals showed greater separation among sequence strategies, with more significant differences involving both repetition and alternation relative to the other patterns (**Table 1**). This separation was further evident in OE baseline animals, where both repetition and alternation occurred significantly more frequently than the remaining strategies, while the two predominant strategies did not differ significantly from each other (**Table 2**). In OE Aβ42 animals, the distribution shifted further toward repetition: LLLL/RRRR remained significantly higher than each of the other strategies, including LRLR/RLRL, whereas the significance of LRLR/RLRL relative to several other strategies was reduced (**Table 3**).

**Figure 4.**
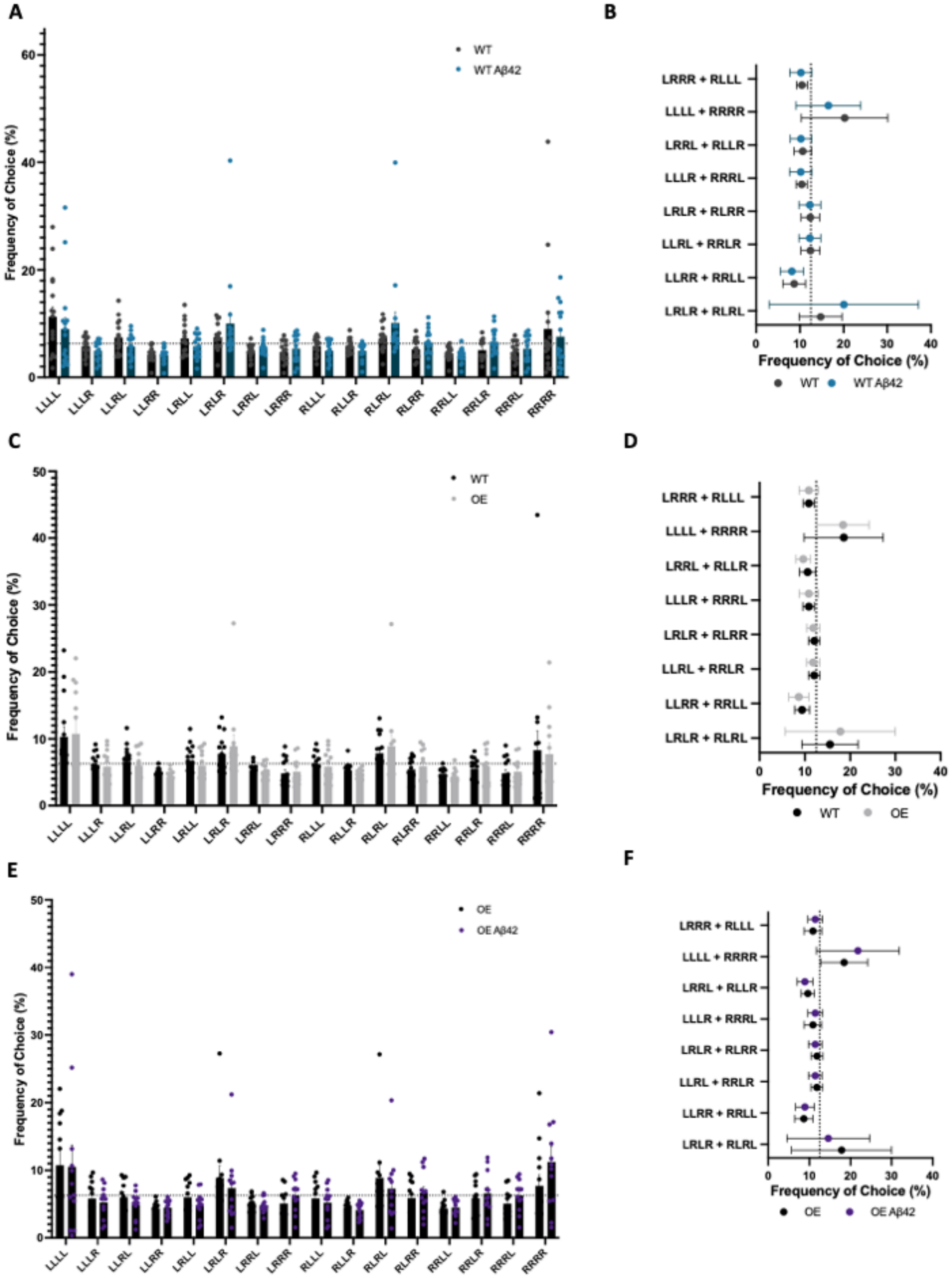
Spontaneous turning strategies in the free-movement pattern Y-maze. Left and right turns recorded over 60 min were grouped into overlapping four-turn sequences. Panels **A** and **B** compare uninjected WT and Aβ42-injected WT fish (n = 16 per group). Panels **C** and **D** compare uninjected WT (n = 14) and uninjected OE fish (n = 12). Panels **E** and **F** compare uninjected OE and Aβ42-injected OE fish (n = 12 per group). Panels A, C, and E show the frequencies of all 16 sequences. Panels B, D, and F show the frequencies of eight complementary sequence pairs. LLLL/RRRR denotes repetition and LRLR/RLRL denotes alternation. Reference lines indicate the expected frequencies for individual sequences (6.25%) and complementary pairs (12.5%) under independent, equally probable left and right turns. Points represent individual fish; bars show group means with SEM.

**Table 1.**
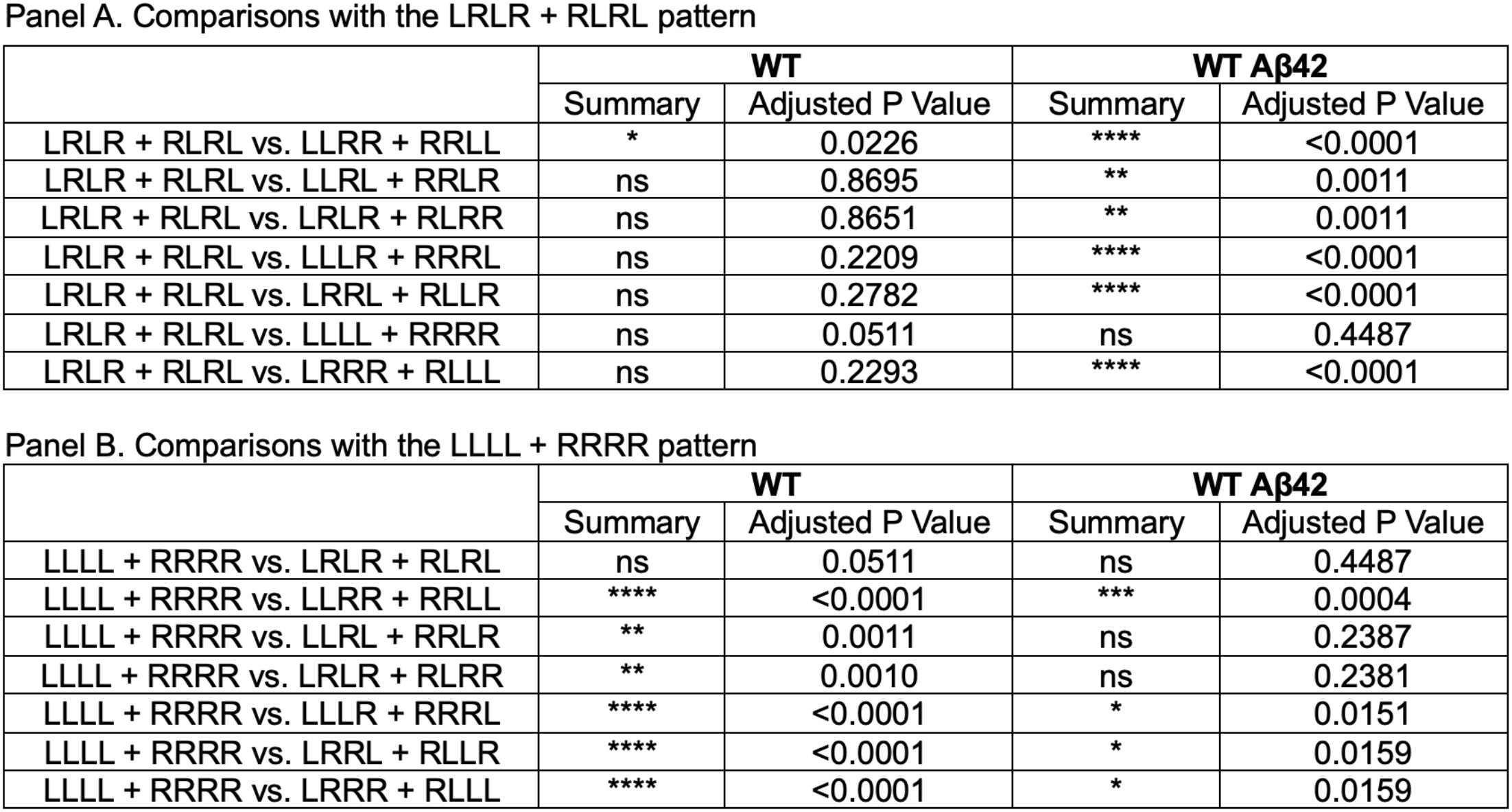
Šídák’s multiple comparisons of paired tetragram strategies in WT and WT Aβ42 fish.

| Panel A. Comparisons with the LRLR + RLRL pattern |  |  |  |  |
|  | **WT** |  | **WT Aβ42** |  |
|  | Summary | Adjusted P Value | Summary | Adjusted P Value |
|---|---|---|---|---|
| LRLR + RLRL vs. LLRR + RRLL | * | 0.0226 | **** | <0.0001 |
| LRLR + RLRL vs. LLRL + RRLR | ns | 0.8695 | ** | 0.0011 |
| LRLR + RLRL vs. LRLR + RLRR | ns | 0.8651 | ** | 0.0011 |
| LRLR + RLRL vs. LLLR + RRRL | ns | 0.2209 | **** | <0.0001 |
| LRLR + RLRL vs. LRRL + RLLR | ns | 0.2782 | **** | <0.0001 |
| LRLR + RLRL vs. LLLL + RRRR | ns | 0.0511 | ns | 0.4487 |
| LRLR + RLRL vs. LRRR + RLLL | ns | 0.2293 | **** | <0.0001 |
| Panel B. Comparisons with the LLLL + RRRR pattern |  |  |  |  |
|  | **WT** |  | **WT Aβ42** |  |
|  | Summary | Adjusted P Value | Summary | Adjusted P Value |
| LLLL + RRRR vs. LRLR + RLRL | ns | 0.0511 | ns | 0.4487 |
| LLLL + RRRR vs. LLRR + RRLL | **** | <0.0001 | *** | 0.0004 |
| LLLL + RRRR vs. LLRL + RRLR | ** | 0.0011 | ns | 0.2387 |
| LLLL + RRRR vs. LRLR + RLRR | ** | 0.0010 | ns | 0.2381 |
| LLLL + RRRR vs. LLLR + RRRL | **** | <0.0001 | * | 0.0151 |
| LLLL + RRRR vs. LRRL + RLLR | **** | <0.0001 | * | 0.0159 |
| LLLL + RRRR vs. LRRR + RLLL | **** | <0.0001 | * | 0.0159 |

**Table 2.**
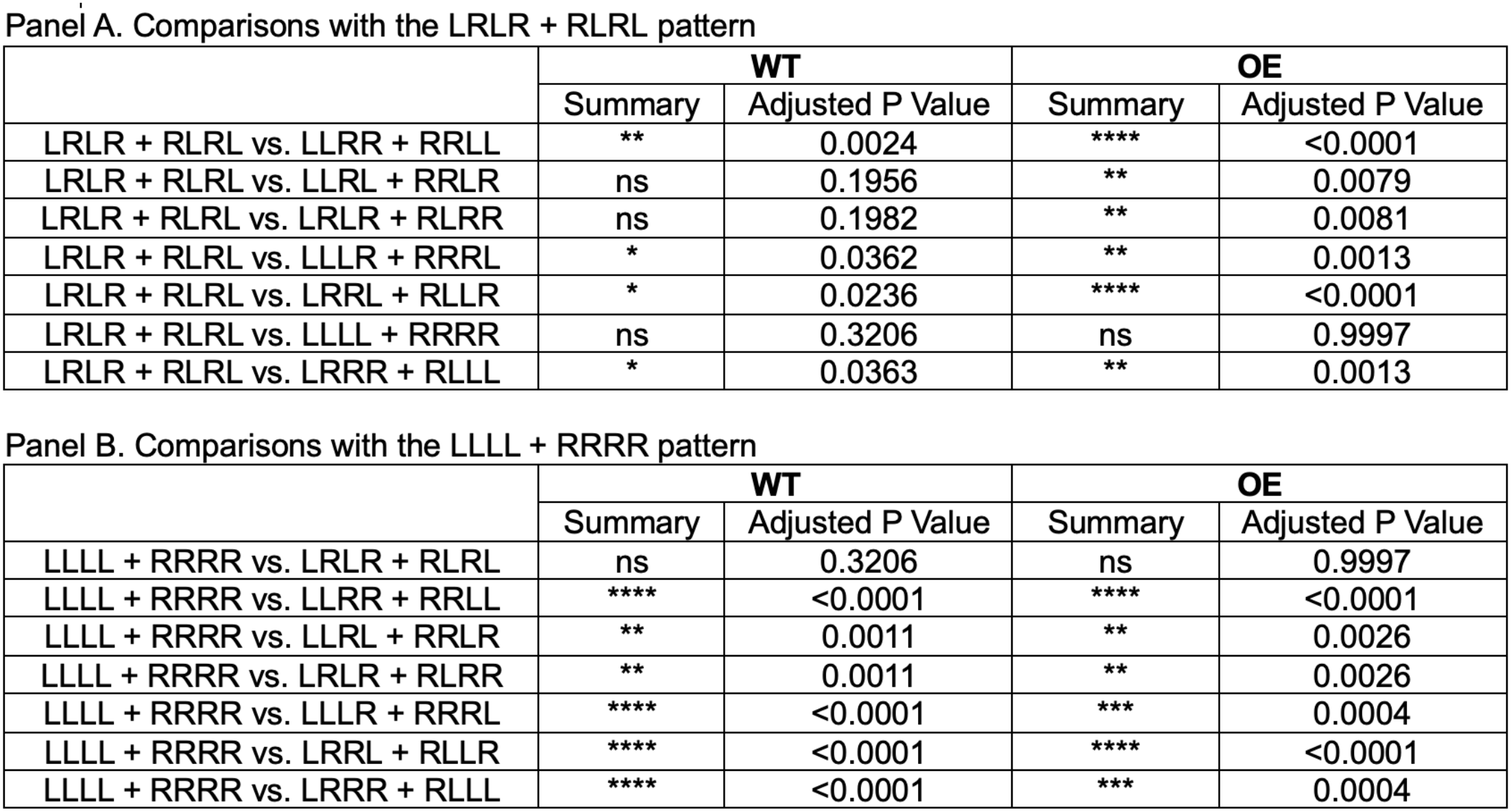
Dunnett’s multiple comparisons of paired tetragram strategies in WT and OE fish.

| Panel A. Comparisons with the LRLR + RLRL pattern |  |  |  |  |
|  | WT |  | OE |  |
|  | Summary | Adjusted P Value | Summary | Adjusted P Value |
|---|---|---|---|---|
| LRLR + RLRL vs. LLRR + RRLL | ** | 0.0024 | **** | <0.0001 |
| LRLR + RLRL vs. LLRL + RRLR | ns | 0.1956 | ** | 0.0079 |
| LRLR + RLRL vs. LRLR + RLRR | ns | 0.1982 | ** | 0.0081 |
| LRLR + RLRL vs. LLLR + RRRL | * | 0.0362 | ** | 0.0013 |
| LRLR + RLRL vs. LRRL + RLLR | * | 0.0236 | **** | <0.0001 |
| LRLR + RLRL vs. LLLL + RRRR | ns | 0.3206 | ns | 0.9997 |
| LRLR + RLRL vs. LRRR + RLLL | * | 0.0363 | ** | 0.0013 |
| Panel B. Comparisons with the LLLL + RRRR pattern |  |  |  |  |
|  | WT |  | OE |  |
|  | Summary | Adjusted P Value | Summary | Adjusted P Value |
| LLLL + RRRR vs. LRLR + RLRL | ns | 0.3206 | ns | 0.9997 |
| LLLL + RRRR vs. LLRR + RRLL | **** | <0.0001 | **** | <0.0001 |
| LLLL + RRRR vs. LLRL + RRLR | ** | 0.0011 | ** | 0.0026 |
| LLLL + RRRR vs. LRLR + RLRR | ** | 0.0011 | ** | 0.0026 |
| LLLL + RRRR vs. LLLR + RRRL | **** | <0.0001 | *** | 0.0004 |
| LLLL + RRRR vs. LRRL + RLLR | **** | <0.0001 | **** | <0.0001 |
| LLLL + RRRR vs. LRRR + RLLL | **** | <0.0001 | *** | 0.0004 |

**Table 3.**
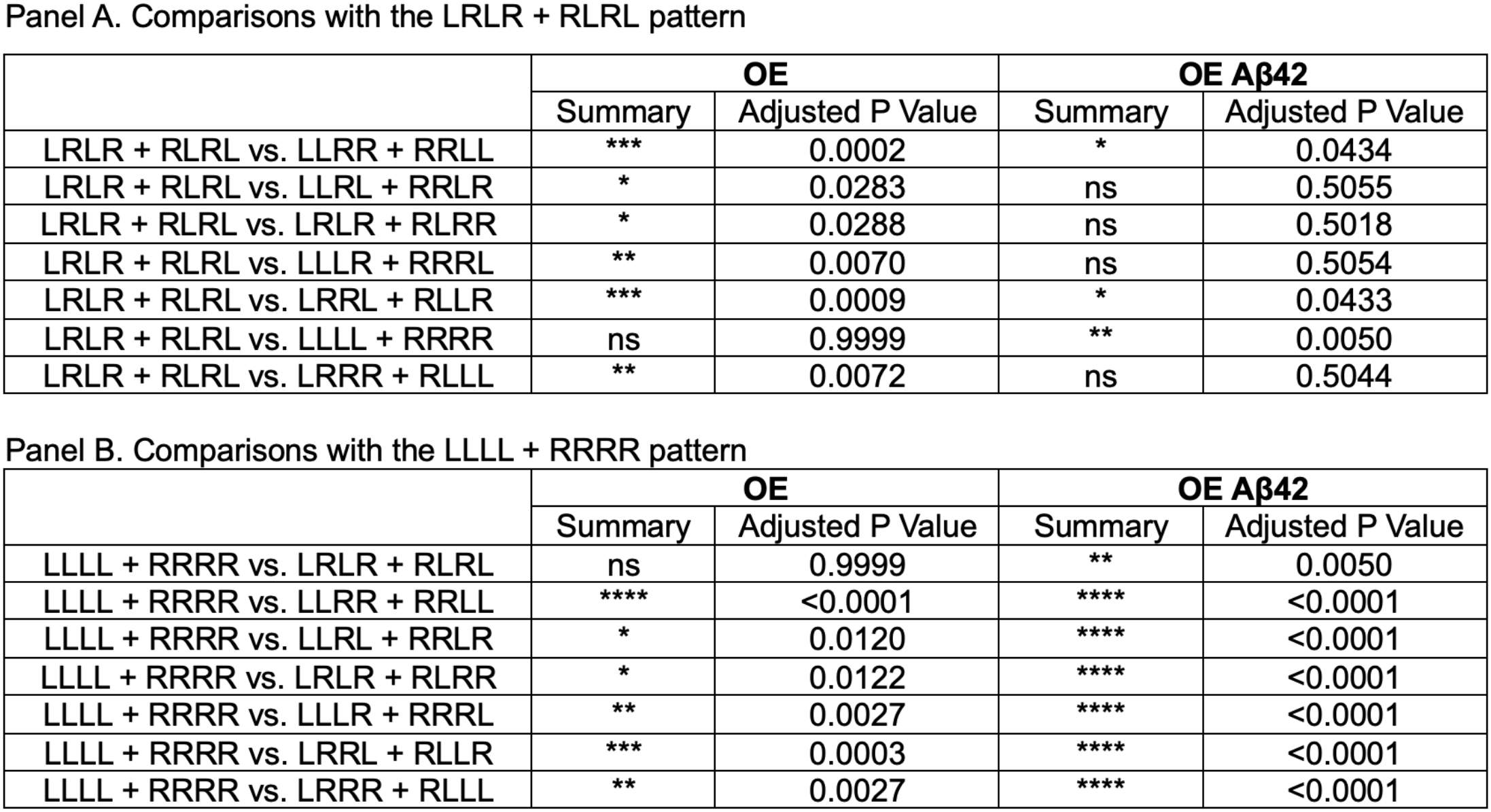
Dunnett’s multiple comparisons of paired tetragram strategies in OE and OE Aβ42 fish.

| Panel A. Comparisons with the LRLR + RLRL pattern |  |  |  |  |
|  | OE |  | OE Aβ42 |  |
|  | Summary | Adjusted P Value | Summary | Adjusted P Value |
|---|---|---|---|---|
| LRLR + RLRL vs. LLRR + RRLL | *** | 0.0002 | * | 0.0434 |
| LRLR + RLRL vs. LLRL + RRLR | * | 0.0283 | ns | 0.5055 |
| LRLR + RLRL vs. LRLR + RLRR | * | 0.0288 | ns | 0.5018 |
| LRLR + RLRL vs. LLLR + RRRL | ** | 0.0070 | ns | 0.5054 |
| LRLR + RLRL vs. LRRL + RLLR | *** | 0.0009 | * | 0.0433 |
| LRLR + RLRL vs. LLLL + RRRR | ns | 0.9999 | ** | 0.0050 |
| LRLR + RLRL vs. LRRR + RLLL | ** | 0.0072 | ns | 0.5044 |
| Panel B. Comparisons with the LLLL + RRRR pattern |  |  |  |  |
|  | OE |  | OE Aβ42 |  |
|  | Summary | Adjusted P Value | Summary | Adjusted P Value |
| LLLL + RRRR vs. LRLR + RLRL | ns | 0.9999 | ** | 0.0050 |
| LLLL + RRRR vs. LLRR + RRLL | **** | <0.0001 | **** | <0.0001 |
| LLLL + RRRR vs. LLRL + RRLR | * | 0.0120 | **** | <0.0001 |
| LLLL + RRRR vs. LRLR + RLRR | * | 0.0122 | **** | <0.0001 |
| LLLL + RRRR vs. LLLR + RRRL | ** | 0.0027 | **** | <0.0001 |
| LLLL + RRRR vs. LRRL + RLLR | *** | 0.0003 | **** | <0.0001 |
| LLLL + RRRR vs. LRRR + RLLL | ** | 0.0027 | **** | <0.0001 |

## Discussion

Our findings indicate that sustained human FMNL2 overexpression is associated with histological and behavioral changes in adult zebrafish, including reduced BLBP-ZO1 overlap, lower normalized SV2 counts, increased microglial counts, and altered novel-tank exploration. Several histological changes were also observed following acute Aβ42 exposure in wild-type animals, whereas additional Aβ42 exposure produced no statistically detectable change in these endpoints within the overexpression background. These observations extend previous work identifying FMNL2 as a regulator of amyloid-associated gliovascular remodeling and suggest that persistent FMNL2 elevation can influence brain phenotypes even without an acute amyloid challenge. One possible interpretation is that the consequences of FMNL2 elevation depend on its duration and biological context. Previous work showed that FMNL2 is required for gliovascular remodeling following amyloid exposure, supporting its involvement in an acute response to pathology. A transient reduction in astroglial-vascular association may facilitate remodeling and the handling of extracellular material. Persistent FMNL2 elevation, however, could maintain an altered glial-vascular organization beyond the period in which that response is beneficial. The reduced BLBP-ZO1 overlap, lower SV2 counts, and increased microglial counts observed in OE animals are consistent with this possibility. Nevertheless, the present study does not directly demonstrate a transition from an initially protective response to a detrimental state, because it does not track these processes over time or manipulate the duration of FMNL2 expression.

The absence of a statistically detectable additional histological response to Aβ42 in OE animals warrants further consideration. Chronic FMNL2 elevation may engage processes that overlap with those recruited by acute amyloid exposure, producing convergence in the measured endpoints. Alternatively, baseline changes in OE animals may constrain the range over which these markers can change, or additional responses may occur at times or through pathways not captured by the present measurements. Limited biological replication may also reduce sensitivity to further changes. Thus, the findings are compatible with an altered response to amyloid in the OE background, but they do not establish that the response is saturated or that the brain has lost the capacity to respond to additional challenges. Distinguishing these possibilities will require direct assessment of genotype-treatment interactions, higher effect sizes, and responses across a broader temporal and exposure range.

These observations are relevant to Alzheimer’s disease, in which elevated FMNL2 expression has been associated with cerebrovascular pathology and amyloid deposition ^24,25^. The histological changes observed in FMNL2-overexpressing zebrafish without an acute Aβ42 challenge raise the possibility that elevated FMNL2 contributes to disease-associated cellular changes rather than serving solely as a marker of an ongoing response. In this model, amyloid-induced FMNL2 elevation could participate in remodeling that becomes persistent when the pathological stimulus is not resolved. Sustained changes in glial-vascular organization, synaptic markers, and microglial state could then contribute to the maintenance of an altered tissue environment. A reciprocal relationship between FMNL2 elevation and amyloid-associated pathology is therefore plausible. Amyloid may increase FMNL2 expression, while persistent FMNL2 activity could influence processes involved in extracellular aggregate handling and tissue homeostasis. Alternatively, FMNL2 elevation could precede amyloid accumulation and alter the conditions under which pathology develops. The present experiments do not establish the direction of this relationship, because they do not measure amyloid clearance, longitudinal amyloid accumulation, or the temporal sequence of endogenous FMNL2 elevation. These possibilities should therefore be considered testable models rather than demonstrated feedback mechanisms. The association of FMNL2 with cardiovascular risk and cerebrovascular pathology provides an additional reason to investigate these relationships. Altered FMNL2-dependent remodeling may be one process through which vascular and amyloid-related stresses converge on the glial-vascular interface. However, the current study does not directly test cardiovascular risk factors, vascular injury, or cerebral perfusion. Its contribution is to show that sustained FMNL2 overexpression is associated with changes in readouts relevant to this interface, providing a basis for examining how FMNL2 influences responses to combined vascular and amyloid-related stress.

The behavioral findings suggest that chronic FMNL2 elevation is accompanied by changes extending beyond the histological endpoints examined. In the novel tank test, OE animals showed a higher composite boldness index than WT controls, together with differences in swimming speed at selected time points. Greater exploration of the upper tank may be consistent with altered novelty responses or risk-related behavior. However, the composite index incorporates both vertical position and distance traveled, so increased locomotor activity may contribute to the higher score. The findings are therefore best interpreted as an altered exploratory profile rather than a specific demonstration of impaired risk assessment. Additional Aβ42 exposure did not produce a statistically detectable change in the composite index within the OE comparison. The Y-maze findings provide a complementary description of spontaneous exploratory organization. Repetition and alternation were prominent turning strategies, and their relative use varied descriptively across the experimental comparisons. OE animals exposed to Aβ42 showed a stronger preference for repetition relative to alternation, raising the possibility that combined FMNL2 elevation and amyloid exposure influence the organization of sequential choices. However, significant differences among strategies within a group do not independently establish that strategy use differs between groups. Direct between-group analyses are needed before attributing these distributions to FMNL2 overexpression or Aβ42 treatment. Temporal organization may provide further insight into how animals explore the maze. Nevertheless, our results identify aspects of exploratory behavior that can be examined in future studies using adequately powered longitudinal analyses and complementary cognitive tasks. Taken together, the behavioral data support further investigation of FMNL2-associated changes in exploration, while remaining insufficient to establish a specific deficit in working memory or executive function.

Several limitations define the scope of these conclusions. First, constitutive transgenic expression does not distinguish the effects of FMNL2 during development from those of sustained elevation in the adult brain. An adult-inducible model would help establish whether the observed changes arise after adult FMNL2 elevation and whether they are reversible. Second, the histological measurements provide marker-based assessments. BLBP-ZO1 overlap does not directly measure vascular permeability, SV2 labeling does not establish synaptic function, and L-plastin labeling and morphology do not fully characterize microglial identity or functional state. Complementary structural and functional measurements are therefore needed to determine the physiological consequences of these changes. The present findings should be interpreted as evidence concerning sustained FMNL2 elevation and acute amyloid responses in zebrafish, with their relevance to chronic mammalian disease requiring further investigation.

Several questions emerge based on our findings. A priority is to determine how sustained FMNL2 elevation interacts with persistent amyloid exposure. Combining FMNL2 overexpression with a chronic amyloid model would help establish whether the overlapping histological changes observed here remain stable, progress, or diverge as pathology develops. Measurements of amyloid burden and clearance alongside glial-vascular organization, vascular permeability, synaptic function, and microglial responses would distinguish changes in marker expression from changes in tissue function. The timing and regulation of FMNL2 elevation also require investigation. Longitudinal studies could determine whether increased expression precedes amyloid accumulation, follows vascular or amyloid-related injury, or participates in a reciprocal process. Identifying the signals that induce FMNL2 in different cell types would clarify whether its regulation is specific to amyloid or reflects a broader response to tissue stress. Adult-inducible and cell-type-restricted approaches would further help separate developmental effects from adult responses and establish which cellular populations contribute to the phenotype. Finally, the downstream mechanisms linking FMNL2-dependent actin regulation to glial-vascular organization remain to be defined. Studies of cytoskeletal remodeling, adhesion complexes, and signaling between glial and endothelial cells could identify how elevated FMNL2 changes these interactions. Restoring FMNL2 toward baseline or manipulating its downstream effectors would test whether the observed changes are reversible and dependent on continued FMNL2 elevation. Such experiments would provide a direct way to evaluate the proposed distinction between an acute remodeling response and the consequences of persistent activation.

The importance of developmental timing is reinforced by a recent preprint reporting that an Fmnl2 mutation disrupting autoinhibition causes brain and vertebral abnormalities in mice ^40^. The mutant protein showed increased actin-regulatory activity despite reduced abundance, whereas Fmnl2 knockout mice lacked the gross anatomical abnormalities. These findings support the broader premise that dysregulated FMNL2 activity can affect nervous-system organization. In our constitutive overexpression model, developmental effects could contribute to the phenotypes measured in adulthood.

In our study, we found that sustained human FMNL2 overexpression in zebrafish is associated with reduced BLBP-ZO1 overlap and SV2 labeling, increased microglial counts, and altered exploratory behavior. Several histological changes resemble those observed after acute Aβ42 exposure in wild-type animals, while additional Aβ42 exposure produces no statistically detectable change in these endpoints within the OE background. These findings support further investigation of persistent FMNL2 elevation as a contributor to altered glial-vascular organization and associated cellular responses. Establishing the temporal, cellular, and functional basis of these effects will clarify their relevance to the intersection of cerebrovascular pathology and Alzheimer’s disease.

## Acknowledgements

We thank Dr. John Copeland for providing the human FMNL2 cDNA used to generate the transgenic zebrafish line. We thank the imaging facility at the Taub Institute for Research on Alzheimer’s Disease and the Aging Brain. We also thank the staff of the Zebrafish Core Facility at Columbia University Irving Medical Center’s Institute of Comparative Medicine for zebrafish care and husbandry. This work was supported by Schaefer Research Scholars Award, the Taub Institute Grants for Emerging Research (TIGER), the Thompson Family Foundation’s TAME-AD program to C.K,, and the Toffler Scholar Program to P.B. Zebrafish maintenance supporting this study was funded in part by the National Institute of NIH/NINDS under award number R21NS149249 to C.K. The content is solely the responsibility of the authors and does not necessarily represent the official views of the National Institutes of Health.

## Author contributions

Conceived the study and defined the research questions: C.L., E.Y., and C.K. Designed the experiments and developed the assays or analysis methods: C.L. and E.Y. Supervised the research: E.Y. and C. K. Funding: C.K. Generated, screened, and validated the FMNL2 transgenic line: H.T. Performed the injections, tissue collection, immunostaining, and microscopy: H.T., P.B., and C.L. Conducted the novel tank and Y-maze experiments: C.L. and E.Y. Wrote the image-processing and behavioral-analysis code: C.L. Quantified histological outcomes and performed statistical analyses: C.L. and E.Y. Analyzed behavioral recordings and performed statistical analyses: C.L. and E.Y. Organized the raw measurements, metadata, and supplementary datasets. C.L. Prepared the figures and tables: C.L. Writing – original draft: C.L., E.Y., P.B. and C.K. Writing – review and editing: C.L., E.Y., H.T., P.B., and C.K. All authors read and approved the final manuscript.

## Supplementary files

**Supplementary File 1.** Python codes used for image analysis.

**Supplementary File 2.** Python codes used for behavioral analysis.

**Supplementary Figure 1.** PCR-based genotyping of overexpression lines. Representative genotyping gel showing the expected 809-bp PCR amplicon. The expected band is outlined by a red rectangle.

**Supplementary Table 1.** Supporting parameters for classification of microglia at baseline conditions.

**Supplementary Table 2.** Supporting parameters for classification of microglia at amyloid treatment conditions.

